# Genetic controls of mouse *Tas1r3*-independent sucrose intake

**DOI:** 10.1101/379347

**Authors:** Cailu Lin, Michael G. Tordoff, Xia Li, Natalia P. Bosak, Masashi Inoue, Yutaka Ishiwatari, Gary K. Beauchamp, Alexander A. Bachmanov, Danielle R. Reed

## Abstract

We have previously shown that variation in sucrose intake among inbred mouse strains is due in part to polymorphisms in the *Tas1r3* gene, which encodes a sweet taste receptor subunit and accounts for the *Sac* locus on distal Chr4. To discover other quantitative trait loci (QTLs) influencing sucrose intake, voluntary daily sucrose intake was measured in an F^2^ intercross with the *Sac* locus fixed; in backcross, reciprocal consomic strains; and in single- and double-congenic strains. Chromosome mapping identified *Scon3*, located on Chr9, and epistasis of *Scon3* with *Scon4* on Chr1. Mice with different combinations of *Scon3* and *Scon4* genotypes differed more than threefold in sucrose intake. To understand how these two QTLs influenced sucrose intake, we measured resting metabolism, glucose and insulin tolerance, and peripheral taste responsiveness in congenic mice. We found that the combinations of *Scon3* and *Scon4* genotypes influenced thermogenesis and the oxidation of fat and carbohydrate. Results of glucose and insulin tolerance tests, peripheral taste tests, and gustatory nerve recordings ruled out plasma glucose homoeostasis and peripheral taste sensitivity as major contributors to the differences in voluntary sucrose consumption. Our results provide evidence that these two novel QTLs influence mouse-to-mouse variation in sucrose intake and that both likely act through a common postoral mechanism.

## Introduction

Mice and rats avidly drink sucrose solutions, and they often become obese within a few weeks if provided with a sucrose solution in addition to food (Ramirez 1987, Sclafani 1987). Understanding the controls of sucrose consumption is thus of theoretical and practical importance for the study of motivated behavior and of diseases associated with obesity. However, the overconsumption of sucrose and ensuing obesity differ by the genetic makeup of the rodent strain (Glendinning, Breinager et al. 2010). These differences may be due in part to how much sucrose the mouse or rat drinks and in part to its metabolism. Here we focus on the genetics of sucrose consumption, which is highly preferred by nearly all rodents and is usually at least equally or perhaps more potent than some other nutritive sugars, such as glucose, in stimulating intake and weight gain (Kanarek and Orthen-Gambill 1982, Sclafani and Mann 1987).

A genetic component to sucrose consumption is supported by marked differences in the avidity for sucrose among inbred mouse and rat strains (Lush 1989, Lewis, Ahmed et al. 2005, Tordoff, Alarcon et al. 2008); moreover, rat strains have been selectively bred to drink large volumes of solutions containing sugar and other sweeteners (Dess 2000). In laboratory mice, the *Sac* (saccharin preference) locus influences intake of sweeteners, including sucrose (Fuller 1974). We and others have identified *Tas1r3*, the gene underlying the *Sac* locus, which codes for a G-protein-coupled receptor (Bachmanov, Li et al. 2001, Kitagawa, Kusakabe et al. 2001, Max, Shanker et al. 2001, Montmayeur, Liberles et al. 2001, Sainz, Korley et al. 2001) that combines with the T1R2 subunit to form the T1R2+T1R3 sweet taste receptor (Nelson, Hoon et al. 2001).

Variation in the T1R2+T1R3 sweet receptor accounts for differences in sweetener intake among several mouse strains (Reed, Li et al. 2004). Indeed, it plays such a large role in determining sweetener consumption (Inoue, Reed et al. 2004, Reed, Li et al. 2004) that it may overshadow contribution of other genetic loci, existence of which we hypothesized based on several pieces of evidence. First, strains of mice with the same *Tas1r3* haplotype show substantial variation in sweetener consumption, with a large combined effect of other loci (Reed, Li et al. 2004). Second, *Tas1r3* does not account for variation in sweetener intake in rats, and thus other genes must be responsible for this variation (Lu, McDaniel et al. 2005). We have confirmed this hypothesis by performing a genome scan that identified several additional quantitative trait loci (QTLs) for sucrose consumption (see the companion paper (Lin 2020)). In addition to the *Sac/Tas1r3* locus (named according to nomenclature as <u>**S**</u>ucrose <u>**con**</u>sumption, QTL **2**, or *Scon2*), an especially prominent new QTL (*Scon3)* was mapped to chromosome (Chr) 9, and epistatic interaction of *Scon3* with *Scon4* (on Chr1) was detected. When we present the data with chromosome order, *Scon4* (on Chr1) is first and then *Scon3* (on Chr9), especially for the description of genotypes for the epistatic QTLs in Figures. We named the newly detected sucrose QTL on Chr14 as *Scon10* (**S**ucrose **con**sumption, QTL **10**).

Although *Scon2* (*Sac/Tas1r3*), *Scon3* and *Scon4* all influence sucrose intake, the patterns of their phenotypical effects differ: whereas the effects of *Scon2* were most apparent when mice were offered low concentrations of sucrose solutions, the *Scon3* and *Scon4* loci were most apparent when mice were offered high concentrations of sucrose. Thus, to isolate the *Scon3* and *Scon4* loci, we tested mice with higher concentrations of sucrose and designed experiments to reduce or eliminate the effects of the *Scon2* locus. To this end, we adopted a strategy used in immunology to identify histocompatibility genes beyond the major histocompatibility complex (MHC). The approach was to keep the MHC haplotype constant among different strains of inbred mice while allowing other genes to vary, which revealed more histocompatibility antigens and had important practical implications for human transplantation biology (Almoguera, Shaked et al. 2014). Thus, we bred mice that had the same genotype at the *Scon2* locus, which allowed us to uncover non-*Scon2* loci regulating sucrose intake. Specifically, we bred a mouse strain that was congenic at the *Scon2* locus. We intercrossed with an inbred strain that was identical to the congenic line at the *Scon2* locus but differed elsewhere in the genome, and we then conducted a genome-wide screen of these intercrossed mice. The newly discovered *Scon3* and *Scon4* loci were further pursued using consomic and congenic mapping techniques, capturing them in fragments of the donor genome and reducing their size over successive cycles of breeding.

To understand the underlying physiological mechanisms responsible for the different intakes of sucrose by mice of different genotypes, we assessed the congenic mice for their metabolism (with indirect calorimetry), gustatory nerve responses (with electrophysiology), and blood glucose and insulin responses. Our goal was to pinpoint the regions affecting sucrose consumption and gain information that would allow us to narrow the list of potential genes.

## Methods

### Animal husbandry

All animal study procedures were approved by the Monell Chemical Senses Center Institutional Care and Use Committee. Inbred C57BL/6ByJ (B6; stock no. 001139) and 129P3/J (129; stock no. 000690) mice used as progenitor strains for the initial stages of breeding were purchased from The Jackson Laboratory (Bar Harbor, ME, USA); we bred all other mice in the animal vivarium at the Monell Center, located in Philadelphia, Pennsylvania (USA). We fed all mice Rodent Diet 8604 (Harlan Teklad, Madison, WI, USA) and gave them tap water to drink from glass bottles with stainless steel spouts (except while being tested; see below). All mice were housed in a temperature-controlled vivarium at 23°C on a 12:12-hr light cycle, with lights off at 7 pm, barring unusual circumstances (e.g., power outages). Pups were weaned at 21-30 days of age, and they were housed in same-sex groups.

### Breeding overview

We bred and studied four types of mouse mapping populations, all derived from B6 and 129 inbred strains (S1 Table): F_2_ intercross, backcross, consomic, and congenic (single and double) strains. The specific breeding strategy was different for each population, but all were part of a large-scale program to map taste genes (Lin, Fesi et al. 2015, Lin, Fesi et al. 2017).

#### F_2_ intercross

The *F*_*2*_ intercross was designed so that all mice had the B6 allele of the *Tas1r3* gene (Reed, Li et al. 2004). Females of the B6 inbred strain were mated with two males from an incipient congenic strain 129P3/J.C57BL/6ByJ-*Tas1r3* (N_4_F_4_) (Bachmanov, Li et al. 2001, Li, Inoue et al. 2001) to produce F_1_ and then F_2_ generations.

#### Backcross and consomic

We generated serial backcross generations (S1 Table) by intercrossing inbred B6 and 129 strains to produce F_1_ hybrids, and then crossing F_1_ (females) with males from B6 or 129 strain to breed two reciprocal N_2_ hybrids. Starting with N_3_, the selected breeders from the 129.B6 strain were mated with 129 females. We pooled four backcross generations to form one mapping population (N_3_ to N_6_) used as segregating generations for QTL mapping, and also to generate consomic mice (S2 Table) using marker-assisted methods (Lin, Fesi et al. 2015, Lin, Fesi et al. 2017, Lin, Fesi et al. 2018).

#### Congenic strains

We branched all congenic strains from consomic strains by mating heterozygous consomic or partially consomic mice with inbred 129 partners. This strategy allowed us to compare phenotypes of littermates with one copy of the donor region (heterozygous; 129/B6) to those without the donor region (homozygous; 129/129). Each congenic mouse was potentially genetically unique (because the donor region could shorten due to meiotic recombination), so we genotyped all congenic mice to define the donor region breakpoints. We named congenic strains with the prefix “C”: for 129.B6-*Scon4*, we named two strains, C1 and C2; for 126.B6-*Scon3*, we named seven strains as C3-C9. All nine strains descended from three progenitors, and the residual genomes of the congenic substrains were tested and calculated (S3 Table). Specifically, we branched *Scon4* congenic mice by backcrossing one partially consomic male from the N_8_ generation (a heterozygous male from 129-Chr1^B6^) to several inbred 129 females. We branched *Scon3* congenic strains by backcrossing several consomic mice (129.B6-Chr9) from the N_7_ generation to 129 females. We bred the double-congenic mice by mating *Scon3* heterozygous females (B6/129; N_7_) with a homozygous *Scon4* male (B6/B6; N_5_F_2_). Thus, for the *Scon4* locus in the double-congenic strain, all offspring were heterozygous (129/B6). For the *Scon3* locus, half the offspring were heterozygous (129/B6) and half were homozygous (129/129).

#### Limits of breeding

We had difficulty producing enough *Scon3* heterozygous or homozygous congenic mice for in-depth phenotyping. We bred *Scon3* congenic strains with shorter and shorter donor regions, but what few mice we could produce had to be used for breeding. These breeding problems, similar to those encountered with the corresponding homozygous consomics (Lin, Fesi et al. 2015), made it difficult to create a homozygous *Scon3* congenic strain. In contrast, *Scon4* congenic mice were fertile and reproduced easily, so we did in-depth taste tests and other phenotyping on *Scon4*, consomic, and F_2_ mice only.

We had no consomic mice available for Chr14, the location of the *Scon10* locus, so we were unable to investigate this *Scon10* locus.

### Genotyping

We extracted and purified genomic DNA from tail tissue either using a sodium hydroxide method (Truett, Heeger et al. 2000) or by proteinase K digestion followed by high-salt precipitation (Gentra/Qiagen, Valencia, CA, USA). We measured the concentration and purity of DNA using a small-volume spectrophotometer (Nanodrop, Wilmington DE, USA). We genotyped these DNA samples using two methods: microsatellites and single-nucleotide polymorphisms (SNPs). For the microsatellites, we used fluorescently labeled primers to amplify genomic DNA by PCR and scanned the products using an ABI 3100 capillary sequencer (Applied Biosystems, Foster City, CA, USA). For the SNPs, we genotyped using primers and fluorescently labeled probes designed to discriminate between alleles, using an ABI Prism 7000 real-time PCR or Step One system (ABI Assay-by-Design, Applied Biosystems, Foster City, CA, USA). All markers and their physical positions are listed in S4 Table.

### Phenotyping

We used mice at least 8 weeks old to measure their taste solution consumption, body composition, metabolism, glucose and insulin tolerance, and gustatory electrophysiology. Each mouse mapping population was phenotyped separately.

#### Taste testing (two-bottle choice tests)

We performed two types of taste tests: a basic screen for sucrose intake and a more in-depth survey using sucrose and other taste solutions (Tordoff 2001). For all tests, mice were 8 or more weeks old. They were individually housed for at least 2 days before beginning these tests and they were trained to drink deionized water from two graduated drinking tubes for at least 2 days. Daily measurements were made in the middle of the light period by reading fluid volume to the nearest 0.1 mL. We measured body weights before and after each test series.

For the **basic screening test**, mice were offered for 96 hr (4 days) one tube of water and one tube of 300 mM sucrose. In later studies they were offered both tubes containing 300 mM sucrose, because we learned from the earlier tests that mice drank nearly 100% of their daily fluid intake from the 300 mM sucrose tube; providing two tubes of sucrose ensured that even the most avid drinkers would not drink it all in a 24-h test (each drinking tube has a maximum volume of ∼25 mL, so two tubes provided up to 50 ml sucrose solution).

For the **in-depth taste tests**, mice were offered for either 96 or 48 hr one tube of deionized water and another tube with a taste solution (described below), switching positions of the two drinking tubes every day to control for side bias (some mice prefer to drink from one side regardless of the contents of the tube). Between each test series, mice usually had two tubes of water for at least 2 days.

Mice from the F_2_ intercross were tested for 96 hr (4 days) with the following taste solutions in the order listed: 30 mM glycine, 30 mM D-phenylalanine, 1.6 and 20 mM saccharin, 50 and 300 mM sucrose, 300 mM monosodium glutamate (MSG), 3% and 10% ethanol, and 50 mM CaCl_2_. All consomic mice, 40 of the *Scon4* homozygous congenic mice, and the inbred control 129 and B6 mice were tested with the following: 30 mM glycine, 300 mM NaCl, 20 mM saccharin, 300 mM sucrose, 300 mM MSG, and 3% and 10% ethanol. We purchased all taste compounds from Sigma Chemical Co. (St. Louis, MO, USA), except the ethanol, which was purchased from Pharmco Products (Brookfield, CT, USA).

All backcross, *Scon3* and *Scon4* heterozygous congenic, and double-congenic mice were tested with the basic sucrose two-bottle choice test. The homozygous *Scon4* congenics (C1) and control 129 inbred mice were tested in four batches (in addition to those 40 mice that were tested with consomic mice) to efficiently test a large number of taste compounds. We tested as follows, for 96 hours: **batch 1**, a concentration series of 10, 30, 100, 300, and 600 mM sucrose (*N*=19 congenic and *N*=10 controls) for 48 hr; **batches 2 and 3**, a concentration series of 10, 30, 100, 300, 600, and 1200 mM glucose and fructose, respectively (*N*=10 mice per genotype group) for 48 hr; **batch 4**, 4.6% soybean oil, 6% cornstarch, and 10.5% maltodextrin (*N*=19 congenics and 10 controls) for 96 hr.

#### Body composition

We measured the body composition of *Scon4* mice (C1) at 8 and 36 weeks of age using magnetic resonance (MR; Bruker minispec LF110 whole-body composition rat and mouse analyzer; Bruker BioSpin Corp., Billerica, MA, USA). After the second MR scan, we also performed body composition analyses by dual-energy x-ray absorptiometry (DEXA; PIXImus II densitometer; GE software, version 2.00; Lunar Corp., Madison, WI, USA) and by necropsy using anatomic landmarks to identify and remove the organs (Cinti 1999, Hayakawa 2001). We weighed the following organs to the nearest 0.01 g: spleen, heart, pancreas, brain, kidney, liver, six “white” adipose depots (pericardial, inguinal, retroperitoneal, subscapular, mesenteric, and gonadal), and a “brown” adipose depot (subscapular). We also measured body length from the base of the teeth to the anus using electronic calipers (Fowler ProMax, Kelley and Kelly Industrial Supply, Syracuse, NY, USA).

#### Metabolism, food and water intake, and activity

We examined whether *Scon4* genotype affects metabolism, food intake, water intake, or activity, assessing the *Scon4* homozygous congenic (C1) and a control group of inbred 129 mice. Using a TSE LabMaster (version 5.0.6; TSE Systems, Inc., Chesterfield, MO, USA), we measured mice at 8 weeks of age and the same mice again at 36 weeks of age. We trained the mice for 3-5 days in cages that mimicked the experimental cages to ensure they learned to eat and drink appropriately from the suspended food and water containers. We transferred the mice to the experimental cages and measured oxygen consumption, carbon dioxide production, food and water intake, and physical activity for 4 consecutive days. We estimated heat production from carbon dioxide and oxygen consumption corrected for lean body mass.

#### Glucose and insulin tolerance tests

We tested homozygous *Scon4* congenic mice (C1) and 129 inbred mice twice (at 8 weeks and again at 36 weeks of age) for changes in blood glucose in response to exogenous glucose (glucose tolerance test; GTT) or insulin administration (insulin tolerance test; ITT) (Murovets, Bachmanov et al. 2015). All mice were tested during the light phase and had unrestricted access to food prior to testing.

For the GTT, we gave the mice glucose (2 g/kg, 10 ml/kg body weight; Sigma-Aldrich St. Louis, MO, USA) either intraperitoneally (IP) (B6/B6 vs 129/129: *N*=20 vs 12) or by intragastric gavage (IG) (B6/B6 vs 129/129: *N*=21 vs 12). For IP administration, the glucose was dissolved in 0.9% saline; for the IG route it was dissolved in deionized water. For the ITT, we injected the mice (B6/B6 vs 129/129: *N*=19 vs 11) with insulin (2 U/kg, IP; insulin, Novo Nordisk A/S, Bagsvaerd, Denmark).

We sampled blood from the tip of the tail, with duplicate measures at each time point (before and after the administration of glucose or insulin) using a One Touch Ultra glucometer (LifeScan, Inc., USA). For the GTT, we sampled blood at 0, 5, 15, 30, 45, 60, 90, and 120 min; for the ITT, we sampled at 0, 5, 30, 60, 90, and 120 min. For the GTT, animals were held in restraint tubes during the blood draws (they were habituated to them before the tests). During the ITT, mice were unrestrained in their home cages.

#### Gustatory electrophysiology

We conducted an electrophysiological experiment with heterozygous *Scon4* (C1) and homozygous *Scon3* congenic mice (C7) and host 129 inbred mice (5 for each group) as described elsewhere (Inoue, Li et al. 2001, Inoue, McCaughey et al. 2001, Inoue, Reed et al. 2004). Mice ranged from 26 to 50 weeks of age. Activity of the whole chorda tympani nerve in response to lingual application of taste solutions was recorded for the following stimuli (in mM, except for ethanol in %): 20 quinine hydrochloride; 100 NaCl; 10 HCl; 1000 glucose; 1000 maltodextrin; 1000 fructose; 100, 300, 1000 sucrose; 2, 6, 20 saccharin; 100, 300, 1000 MSG; 3, 10 ethanol. Each taste solution was washed over the tongue for 30 sec; between taste solution presentations, the tongue was rinsed with deionized water for at least 1 min. The magnitude of the integrated response at 20 sec after stimulus onset was measured and expressed as a proportion of the average of the previous and following responses to 100 mM NH^4^Cl (Inoue, Reed et al. 2004).

### Data analyses

We performed several steps to prepare the data for the main statistical analysis for each mouse population. We checked the distribution of the sucrose intakes for normality within each mapping population using the Lilliefors test (S5 Table), and nonnormal data were transformed (Delignette-Muller). Male and female mice had similar sucrose intake, so we pooled their data. For each mapping population we separately checked the correlations between voluntary sucrose intake and (a) body weight, (b) daily habitual water intake, and (c) age (**S1 Figure**). (Mice that habitually drank the most water also tended to drink the most sucrose, so we used habitual water intake as a covariate in the relevant statistical analyses.) In addition to intake, we also computed preference scores as a ratio of taste solution intake to total fluid intake (taste solution plus water), in percent, although this preference score was an imperfect measure because many mice drink nearly 100% of their fluid intake as 300 mM sucrose when also offered plain water. For all statistical models we used a type 1 (sequential) sum-of-squares model, and when testing for group differences we used Tukey’s HSD tests except as indicated. We computed statistical tests with R (version 3.3.3) and RStudio (version 1.0.136) and graphed the results using either R or Prism 6 (version 6.05; GraphPad Software, La Jolla, CA, USA).

#### F_2_ intercross

We identified genomic regions shared in common by the F_2_ mice that drank the most sucrose using markers from the 19 mouse autosomes and the X chromosome (R package R/QTL, version 1.41 (Broman, Wu et al. 2003)). We estimated genotype probabilities and genotype errors using the *calc*.*genoprob* function, with interval mapping by maximum likelihood estimation (EM algorithm) for the main-effect QTLs using the *scanone* function. We tested the significance of each marker regression against 1000 permutations of the observed data using the n.*perm* function. We defined the confidence intervals as drops of 1 LOD (logarithm of the odds) from the peak expanded to the nearest outside markers using the *lodint* function. We estimated the explained phenotypic variance for main-effect QTLs as 1 – 10^−2^ ^LOD/ *n*^, where *n* is the sample size, using the LOD score from *scanone* (Broman, Wu et al. 2003). We investigated marker pair interactions using the *scantwo* function, evaluating significance against the 1000 permutation tests and confirming the interaction in a general linear model using the single marker and the interaction of marker pairs as fixed factors.

To assess whether markers identified in the analysis also affected the consumption of other taste solutions (30 mM glycine, 300 mM NaCl, 20 mM saccharin, 300 mM sucrose, 300 mM MSG, and 3% and 10% ethanol) tested in the F_2_ population, we analyzed solution intakes in a general linear model using genotype of each locus as a fixed factor. We analyzed preference scores of these solutions using one-way ANOVA followed by post hoc tests. In addition, we computed Pearson correlations among all taste solutions using intakes (adjusted for habitual water intakes) and preference scores separately.

#### Backcrosses

Using a pooled backcross population (N_3_ to N_6_), we conducted a general linear model analysis of sucrose intake with genotype as a fixed factor. For each marker, we calculated (a) the genotype means, (b) the *p*-value test statistic as the negative base 10 logarithm, and (c) the effect size using Cohen’s *D* (Cohen). We report these values, including confidence intervals (defined by 2 units of –log_10_ *p*-value drop), for the peak marker from the mapping population. Statistical thresholds were computed with a Bonferroni correction to an α level of 0.05 for the number of markers [*N*=67; –log(α/*N*) ≈3.13] (Kanarek and Orthen-Gambill 1982).

#### Consomics

Building on the results from the F_2_ intercross, which detected the main-effect QTL on Chr9 (*Scon3*) that has epistatic interaction with the QTL on Chr1 (*Scon4*), we studied the appropriate consomic mice for these chromosomes. For Chr9 we generated two reciprocal consomic strains, whereas for Chr1 we were able to generate only one consomic strain. For these three consomic strains, we analyzed sucrose intakes with a general linear model, treating strain as a fixed factor and using the relevant host inbred strain for comparisons (129-Chr1^B6^ vs inbred 129, 129-Chr 9^B6^ vs inbred 129, and B6-Chr 9^129^ vs inbred B6). We calculated the genotype effect size with Cohen’s *D*, and we analyzed the intakes of taste solutions other than 300 mM sucrose using the same statistical approach as we used for sucrose. We also used a general linear model to evaluate all consomic and host inbred strains with genotype as a fixed factor and habitual water intake as a covariate, followed by post hoc tests. We analyzed preference scores by strain using *t*-tests.

#### Heterozygous congenic analysis

We analyzed the data from congenic mice using the common segment method to find genomic regions shared by strains that have the same trait (Shao, Sinasac et al. 2010). To this end, for all analyses of congenic mice, we used sucrose intake as the outcome measure and the presence or absence of a donor region as a fixed factor, using a proxy marker for the donor region (*rs3708040* for *Scon4* and *rs13480399* for *Scon3*). If the congenic strain differed from the host inbred strain in voluntary sucrose consumption, we concluded the associated donor region contained the causal genetic variant; if not, we concluded otherwise.

#### Homozygous congenic analysis

We conducted taste tests with a concentration series for sucrose, glucose, and fructose and with a single concentration for soybean oil, cornstarch, and maltodextrin. For the concentration series (e.g., fructose), we conducted a two-way mixed-design ANOVA with strain as the fixed factor (congenics vs host inbred 129) and concentration as the repeated measure. For the single concentrations (e.g., soybean oil), we analyzed solution intakes using strain as a fixed factor (congenics vs inbred 129). When informative (no ceiling effect, i.e., toward 100% preference), we analyzed preference scores in the same way. There were no differences by sex, so sex was not included in the mode.

#### Electrophysiology

Chorda tympani responses to individual taste solutions were not normally distributed, so we used nonparametric Mann-Whitney *U*-tests to assess differences between genotype groups (Inoue, Glendinning et al. 2007), and Bonferroni correction was applied for multiple tests for series concentration of each compound.

#### Body composition

We evaluated the effect of *Scon4* genotype on body composition using a two-way ANOVA (genotype x sex) followed by post hoc tests for all three measures of body composition: MR, DEXA, and necropsy. Within each mapping population (F_2_, consomics, and 129.B6-*Scon4* congenics), we also used two-way ANOVAs (genotype x sex) to determine whether mice with genotypes that increased voluntary sucrose intake also gained more body weight during the 4-day sucrose intake tests.

#### Metabolism

Data for each measure from the TSE LabMaster were averaged over the four 24-hr measurement periods. For each individual, parameters were corrected by lean body mass using the algorithm in the software package provided by the manufacturer. We used *t*-tests to compare groups (congenics B6/B6 vs host inbred 129/129).

#### Glucose and insulin tolerance tests

We analyzed data from GTT and ITT using two-way mixed-design ANOVA with genotype as the fixed factor and time of blood collection as the repeated measure.

#### Evaluating genes and variants in the Scon3 and Scon4 region

Using the results of the congenic mapping, we identified specific genomic regions that likely contained the causal variant(s) for each QTL. Within these regions, we identified the known variants and evaluated them in several ways. First, we made a list of all genetic variants using Mouse Genomes Project-Query SNPs, indels, or structural variations (Anonymous 2011), which contains the results of a large-scale genome sequencing project of many inbred mouse strains, including the 129P2/OlaHsd and C57BL/6J strains (Keane, Goodstadt et al. 2011, Yalcin, Wong et al. 2011), which are closely related to the parental inbred strains used in our study (Yang, Wang et al. 2011). From the list of genetic variants we compiled, we identified sequence variants with the potential to cause functional changes using the Sorting Intolerant from Tolerant (SIFT) algorithm (Anonymous 2012). Twelve genetic variants were excluded from this analysis because we could find no unique identifier for the variant in the dbSNP database (build 138) (Anonymous 2016).

Next, using Mouse Genome Data Viewer (Anonymous 2015), we identified all genes residing within the genomic coordinates of *Scon4* and *Scon3* defined in the congenic strains and classified these candidate genes using the PANTHER classification system (Mi, Muruganujan et al. 2013). Finally, using ToppGene suite (Chen, Bardes et al. 2009), we prioritized candidate genes by comparing their functional annotations against a training set that consisted of 332 known carbohydrate metabolism mouse genes from the KEGG database (Kanehisa, Furumichi et al. 2017). We used the Entrez identifiers of the genes (N=248) within the *Scon3* and *Scon4* regions as input for candidate gene prioritization; therefore, only genes with an appropriate identifier were included.

We reasoned that the causal genes from *Scon3* and *Scon4* jointly contribute to the same pathway, so we identified gene pairs that met this criterion (i.e., one gene-pair member was from the *Scon4* region and the other was from the *Scon3* region). We thus assessed the protein-protein associations by searching the protein names in the STRING database (von Mering, Jensen et al. 2005) with multiple active interaction sources that included three types of data: (1) direct experimental evidence of protein-protein interactions; (2) less direct evidence generated by grouping proteins of known function into a metabolic, signaling, or transcriptional pathway; and (3) *in silico* computation techniques for detecting new protein-protein associations. We exported gene pairs for protein-protein interactions with association scores > 0.9, which were computed by benchmarking them against the training set (von Mering, Jensen et al. 2005), and identified those pairs that fit our criterion.

Finally, we compared expression profiling of genes (*N*=248) within the *Scon3* and *Scon4* critical regions at three mouse tissue sites (striatum, brown and white fat) between 129 and B6 inbred strain using publicly available GEO data (Davis and Meltzer 2007), including the closely related strains 129S1/SvImJ and C57Bl/6J. Grouping three samples of each strain 129P3/J and C57BL/6J (GSE7762; (Korostynski, Piechota et al. 2007)), we conducted microarray-based gene expression analysis for mouse striatum with a web tool GEO2R (http://www.ncbi.nlm.nih.gov/geo/geo2r) by calculating the distribution of fluorescence intensity values for the selected samples and then performing a LIMMA (linear model for microarray data analysis) statistical test (Smyth 2005) after applying log transformation to the data. We downloaded bulk RNASeq data [four samples for each of strain (129S1/SvImJ and C57Bl/6J) for each tissue site (mouse brown and white fat); GSE91067] (Raymond E Soccio 2017), and transformed the transcript count data. We created a matrix for the two groups’ data (129 vs B6) and fitted data with the gene-wise glmFit function of edgeR package (Robinson, McCarthy et al. 2010); we conducted likelihood ratio tests for 129 vs B6 for brown and white fat tissue separately. The differentially expressed genes in the two types of gene profiling data sets were reported as one log^2^-fold change (corresponding to a twofold change) with an associated *p*-value (*p*<0.05) corrected for false discovery (FDR) for multiple tests (Benjamini 1995).

## Results

### Identification of main effect and epistatic QTLs

We mapped a highly significant main-effect QTL on Chr9 (*Scon3*) with a strong effect on 300 mM sucrose intake (explaining 37% of the trait variance) and a significant main-effect QTL on Chr14 (*Scon10*) with a weaker effect (*7*% of trait variance); the B6 allele was dominant and increased sucrose consumption in both cases (**Figure 1A**). The peak *Scon3* LOD score was 28 at marker *rs3023231* (106.8 Mb), with a confidence interval of 105.1-108.1 Mb (between *rs13480395* and *rs4227916*). The two-dimensional genome scan detected that *Scon3* epistatically interacts with a new QTL for 300 mM sucrose intake, *Scon4* on Chr1 (**Figure 1B**), with interaction LOD=1.5, and this epistatic interaction was confirmed by general linear model analysis (**Figure 1C**). While the B6 allele of *Scon3* increased sucrose intake, the B6 allele of *Scon4* decreased sucrose intake. The phenotypic difference among *Scon4* genotypes shows up only with 129/129 genotype of *Scon3*.

**Figure 1.**
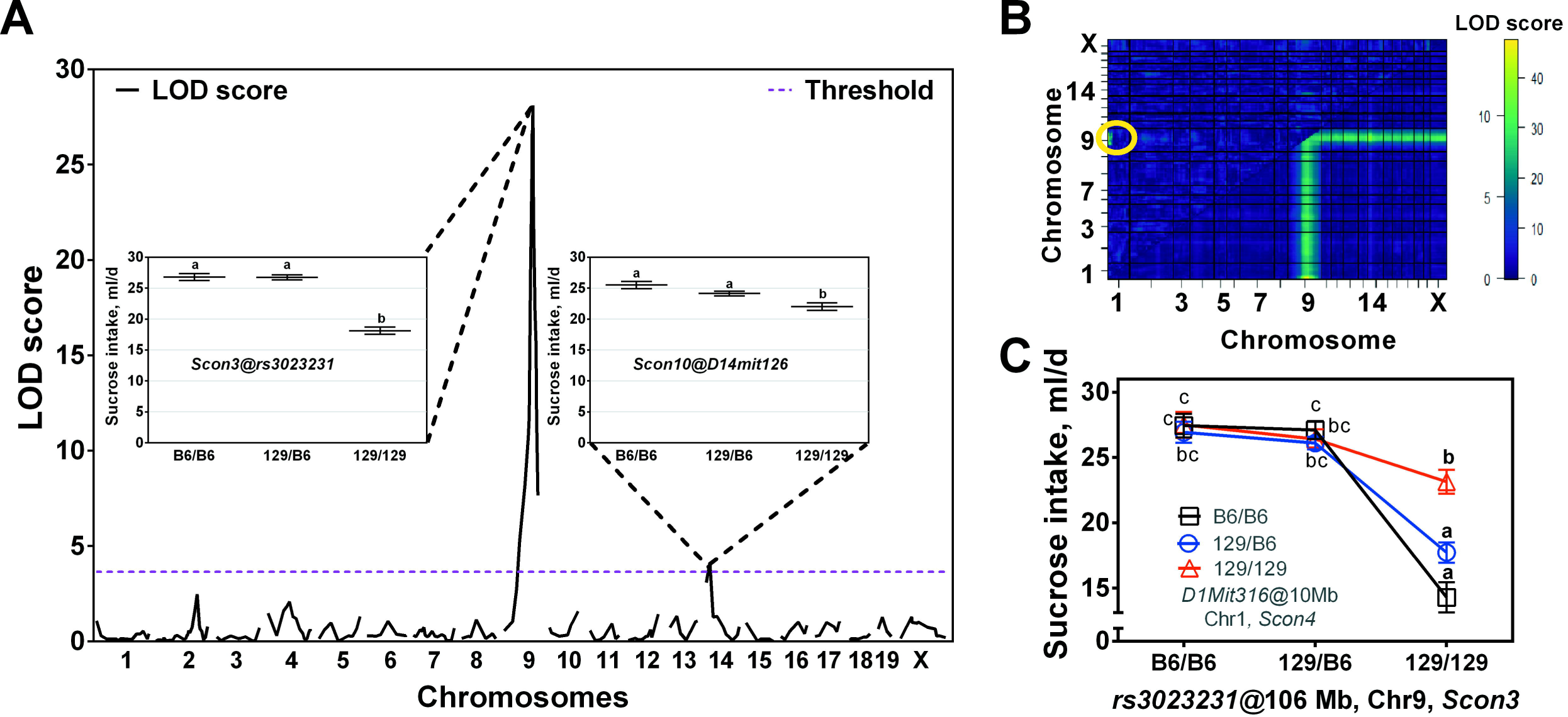
Identification of *Scon3* (a main-effect QTL on Chr9) and *Scon4* (an interacting QTL on Chr1) in C57BL/6ByJ x 129P3/J.C57BL/6ByJ-*Tas1r3* F_2_ mice (N=279) (**A**) A genome-wide scan of main-effect QTLs for 300 mM sucrose intake. The pink horizontal dashed line represents the significant linkage threshold determined by 1000 permutation tests (threshold=3.64). The black curves trace the logarithm of the odds (LOD) scores (y-axis). The genotype effect of the significant loci was confirmed in a general linear model using habitual water intake as a covariate and a post hoc comparison of least-square means (LSM) among the genotypes of these loci tested. Different letters above horizontal lines depicting LSM ± standard error indicate significant differences in daily sucrose consumption by genotype group. (**B**) Heat map for a two-dimensional genome scan with a two-QTL model. The maximum LOD score for the full model (two QTLs plus an interaction) is indicated in the lower right triangle. The maximum LOD score for the interaction model is indicated in the upper left triangle. A color-coded scale displays values for the interaction model (LOD threshold=6.5) and the full model (LOD threshold=9.1) on the left and right, respectively. A yellow circle in the upper left section shows significant interaction (LOD=13.5) between QTLs on *Scon3* and *Scon4*. (**C)** LSM ± standard error for sucrose intakes of the F_2_ mice grouped by genotypes of the markers with the highest epistatic interaction (*D1Mit316* of *Scon4* and *rs3023231 of Scon3*). Different letters indicate statistically significant differences between genotypes (Tukey’s HSD test: *p*<0.05; for details see the Methods section).

Variation at these two loci (*Scon3, Scon4*) had similar effects on the taste preferences of the mice, strongly affecting voluntary consumption of sucrose and a few other taste compounds (**S2 Figure**). For the *Scon4* locus, the only other compounds affected was 20 mM saccharin, but with opposite effect direction: B6 allele of *Scon4* decrease sucrose intake but increase saccharin intake (**S2 Figure**). For the *Scon3* locus, other compounds affected were 3% and 10% ethanol and 300 mM MSG with the same effect direction, as well as 150 mM CaCl^2^ with opposite effect direction (**S2 Figure**). There was no genotype effect (*Scon4* or *Scon3*) on average daily water consumption (**S3 Figure**). Independent of genotype, mice that drank the most 300 mM sucrose drank more MSG and ethanol but not the other taste solutions (**S4 Figure**).

### Confirmation of main effect and epistatic QTLs

#### Backcross

In the backcross population, we detected *Scon3* at the peak marker *rs302323131*; this is the same location as in the F_2_ mapping population and has an allelic effect of the same direction and magnitude, with an –log10 (p vale) of 17.3 and a Cohen’s *D* value of >0.8 (**Figure 2A**). We were not able to test for the effects of *Scon4* because these backcross mice were not genotyped for markers on Chr1.

**Figure 2.**
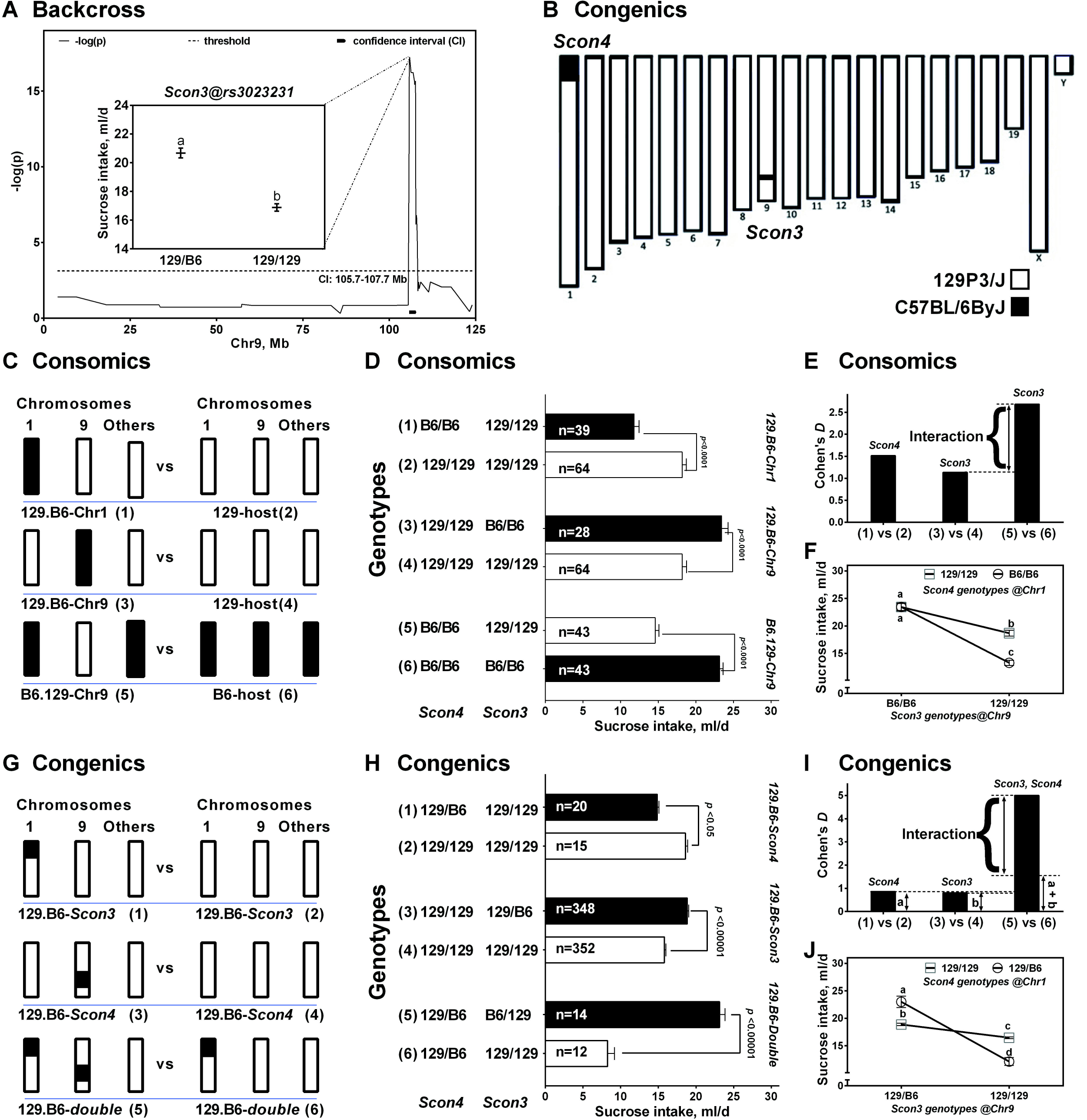
Confirmation of the *Scon3* and *Scon4* loci, and their interaction in backcross (A), consomic (B-E), and congenic (F-J) mice. (**A**) Backcross mice. *Scon3* was detected in the pooled backcross mice (N_3_, N_4_, N_5_, and N_6_; *N*=438). The x-axis is the location of the markers in Mb (see S4 Table) on mouse Chr9; the y-axis is the –log *p*-value of the statistical test by each marker genotype. The small blue bar under the peak indicates the confidence interval (2 units of –log_10_ *p*-value drop from the peak). The horizontal red line shows threshold of statistical significance, calculated using Bonferroni correction [–log(0.05/68) ≈3.13]. The inset shows least-squares means (LSM) ± standard error sucrose intake of mice grouped by peak marker genotype (*rs3023231*). (**B**) A schematic of chromosomes of the 129.B6-double congenic strain, created by mating *Scon4* homozygous congenic males with *Scon3* heterozygous congenic females. These double-congenic mice thus have heterozygous genotypes (B6/129 for all mice) at the *Scon4* locus and segregating genotypes (B6/129 or 129/129) at the *Scon3* locus. (**C-F**) Consomic mice. (**C**) A schematic of the comparisons of mouse consomic strains and their host inbred strains, as well as the corresponding genotypes of *Scon4* and *Scon3* in each mouse strain. (**D**) LSMs and their standard errors for the consomic strains 129.B6-Chr1, 129.B6-Chr9, and B6.129-Chr9, compared with their host inbred strain (129 or B6). (**E**) Effect size in Cohen’s *D* for each group computed by comparing homozygous consomic genotypes vs host inbred genotypes. The large difference in effect sizes between the two reciprocal consomic strains, 129.B6-Chr9 and B6.129-Chr9, supports the epistatic interaction of the homozygous alleles for *Scon3* and *Scon4*: effects of *Scon3* are larger when they are combined with B6/B6 *Scon4* genotype than with 129/129 *Scon4* genotype (compare **Figure 1C**). (**F**) The interaction effect of homozygous alleles between *Scon4* and *Scon3* was confirmed in the consomic and host inbred strains. Different letters indicate statistically significant differences between genotypes. *Scon4*: *F*_(1, 225)_=36.8, *p*<0.00001; *Scon3: F*_(1, 225_)=166.9, *p*<0.00001; *Scon4 × Scon3: F* _(1, 225)_=19.4, *p*<0.0001. (**G-J**) Single- and double-congenic mice. (**G**) Comparisons of littermates from the single-(top two panels) and double-(bottom panel) congenic strains: schematic of chromosomes (left) and corresponding *Scon4* and *Scon3* genotypes (right) in each group of mice. (**H**) LSM and their standard errors for the congenic strains: 129.B6-*Scon4*, 129.B6-*Scon3*, and 129.B6 double congenics, with one copy of B6 donor fragment (B6/129), compared with their littermates without the B6-donor fragment (129/129). (**I**) Effect size in Cohen’s *D* computed for each group of comparisons by comparing heterozygous congenic genotype (B6/129) vs their littermates’ genotype (129/129). A large difference between the effect size observed for the double-congenic strain and the effect sizes for 129.B6-*Scon4* or 129.B6-*Scon3* confirmed the epistatic interaction of the heterozygous alleles between *Scon3* and *Scon4*. (**J**) The interaction effect of heterozygous alleles between *Scon4* and *Scon3* was confirmed in congenic strains: the effects of *Scon3* are larger when they are combined with 129/129 *Scon4* genotype (circles) than with B6/129 *Scon4* genotype (squares; see **Figure 1C**). Different letters indicate statistically significant differences between genotypes.

#### Consomics

The results from the consomics were consistent with those from the F_2_ and backcross populations, confirming the effects of *Scon3* and *Scon4* and their interaction. For *Scon4*, mice with a B6-introgressed Chr1 drank less sucrose solution than did the host control mice (**Figure 2C, D**), with an effect size > 1.5 (**Figure 2E**). For *Scon3*, mice with a B6-introgressed Chr9 drank more sucrose, and mice with the 129-introgressed Chr9 drank less sucrose, compared with host controls, with effect sizes of 1.1 and 2.7, respectively (**Figures 2C, D, E**). Consomic mice with B6 alleles at both *Scon3* and *Scon4* drank much more sucrose than did mice with 129 alleles at both loci (**Figures 2D, F**). We evaluated all consomic and control mice in a single model and found that mice with different genotypes at *Scon4* and *Scon3* differed in voluntary sucrose intake, with a marked increase in mice with certain combinations (e.g., B6/B6 of *Scon4* and B6/B6 *Scon3*) of those alleles (**Figure 2F)**. Consomic mice also differed in their intake of other taste solutions compared with mice from the inbred host strains (**S5 Figure**).

#### 129.B6-Scon4 congenics

The results from congenic mapping support the presence of *Scon4* near the centromere of Chr1 (**Figure 3**). We bred and phenotyped two *Scon4* congenic substrains (C1 and C2; **Figure 3B**; S6 Table). Congenic mice from the C1 substrain (with the donor region from 0 to 38.5 Mb) drank more sucrose than did control mice (**Figure 3C**) and hence retained *Scon4*. However, mice from the C2 congenic strain (with a shorter donor region spanning 24.5-38.5 Mb) drank the same volume of sucrose as did controls (**Figure 3D**) and hence the substrain (C2) did not retain *Scon4*. Comparison of the two substrains defined the *Scon4* critical region as 0-25.4 Mb. This mapping result was supported by the outcome of the general linear model, with the strongest association between sucrose intake and marker genotypes in a 0-25.4 Mb region (**Figure 3A**).

In addition to heterozygous *129*.*B6-Scon4* congenic mice, we produced and phenotyped homozygous congenic mice by intercrossing heterozygotes from substrain C1. *Scon4* congenic homozygotes drank less than one-third the amount of 300 mM sucrose consumed by 129 inbred mice (effect size of 2.6; **Figure 4**), which is consistent with *Scon4* effects in F_2_ (**Figure 1C**).

**Figure 3.**
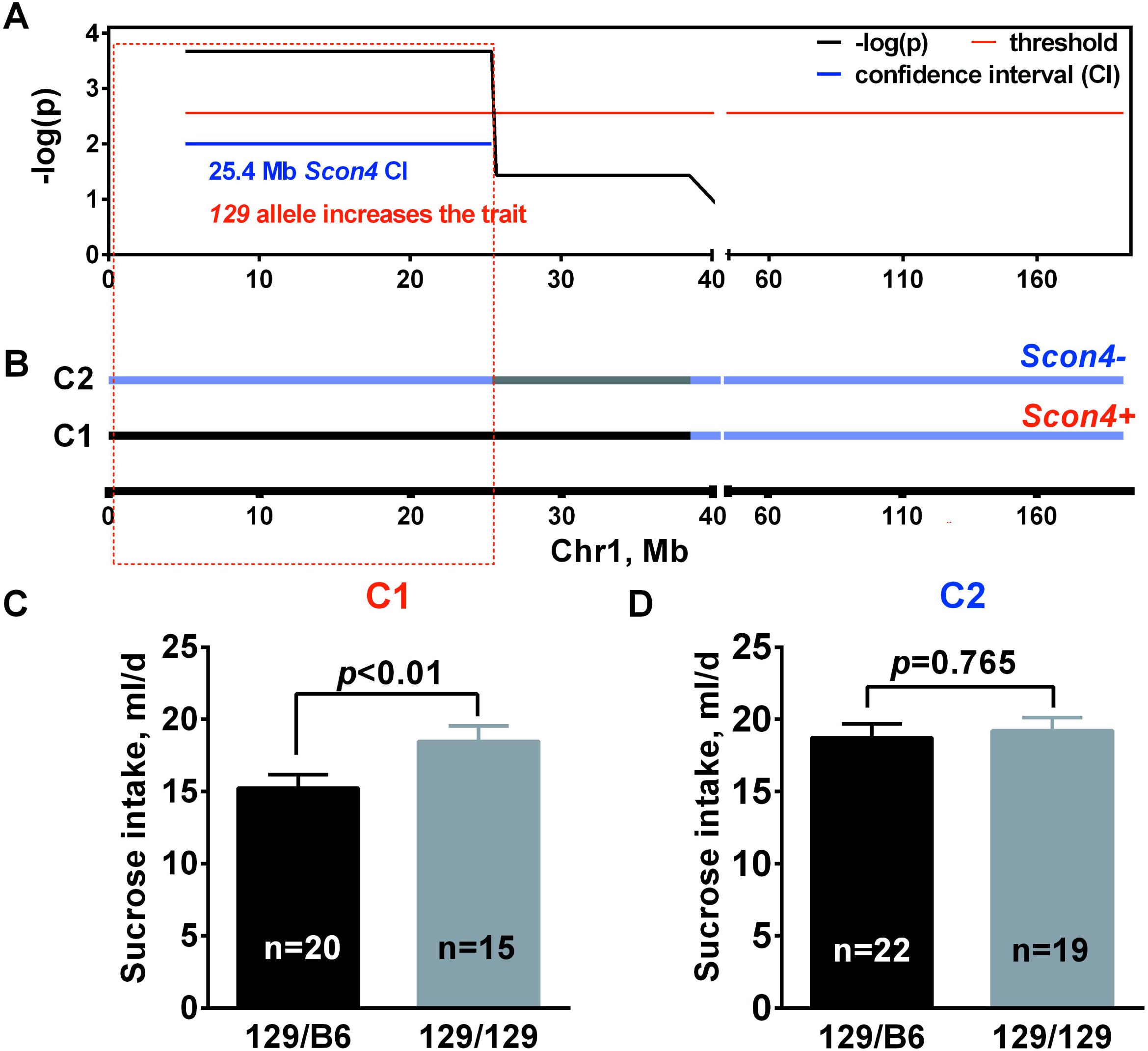
High-resolution mapping of the *Scon4* locus in congenic substrains. Data are from two 129.B6-*Scon4* heterozygous congenic substrains (C1 and C2) with partially overlapping donor fragments (for markers and their positions, see S6 Table). (**A**) Interval mapping of genotype-phenotype associations: using a general linear model, we computed associations between marker genotypes and 300 mM sucrose intake in a combined group including all 129.B6-*Scon4* heterozygous congenic mice (in this analysis, mice were grouped by genotype at each marker rather than by substrain). The x-axis gives marker positions in Mb on Chr1; the y-axis, –log_10_-transformed *p*-values. The horizontal red line shows significance threshold [-log_10_(0.05/*N*), where *N* is number of markers tested]; CI = confidence interval. (**B**) Donor regions of the congenic substrains are indicated by the black bar (for the C1 strain that retained *Scon4*) or gray bar (for the C2 strain that did not retain *Scon4*). Blue bars indicate the regions contributed by the host (129) strain; the *Scon4* critical region (red dashed rectangle; 0-25.4 Mb from centromere to *rs13475771*) is retained in the donor region of the C1 (*Scon4*-positive) strain but not in C2 (*Scon4*-negative) strain. (**C and D**) Intake of 300 mM sucrose by the *Scon4-*positive (C1; **C**) and *Scon4-*negative (C2; **D**) strains. Strain comparisons were made using general linear model.

**Figure 4.**
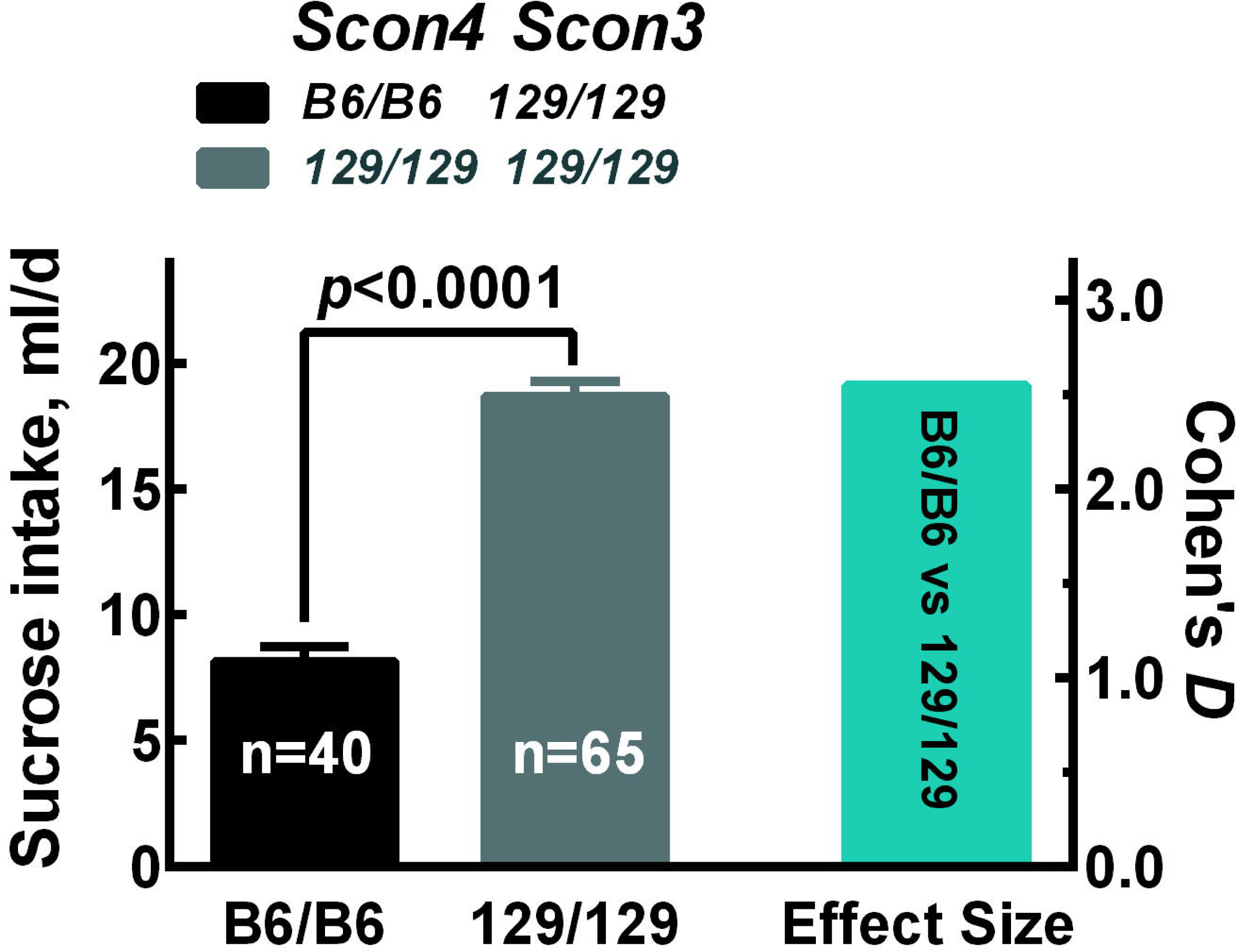
The *Scon4* homozygous allele effect on voluntary sucrose intake. The *Scon4* locus affects 300 mM sucrose consumption in homozygous 129.B6-*Scon4* congenic mice (B6/B6; C1) vs host inbred 129 mice (129/129). Mice with *Scon4* (B6/B6) and *Scon3* (129/129) have lower sucrose intakes than mice with *Scon4* (129/129) and *Scon3* (129/129). The effect size of Cohen’s D is shown in right blue bar.

#### 129.B6-Scon3 congenics

The results from congenic mapping support the presence of *Scon3* in the same region of Chr9 (**Figure 5**) as identified in the F_2_ and backcross mapping populations. We bred and phenotyped seven *Scon3* congenic strains (C3–C9; **Figure 5B**; S7 Table) and found that mice from five strains (C5-C9) with donor fragments including the 1.2 Mb critical region (105.7-106.9 Mb) drank more 300 mM sucrose than did control mice, whereas mice from two strains (C3, C4) without this critical region did not differ from control mice (**Figure 5C**). The mapping result was supported by the outcome of the general linear model, with the strongest relationship between sucrose intake and genotype within this 1.2 Mb region (**Figure 5A**). This region contains 6 noncoding RNA genes, 33 protein-coding genes, 6 pseudogenes, and 9 processed transcripts (**Figure 5D**).

**Figure 5.**
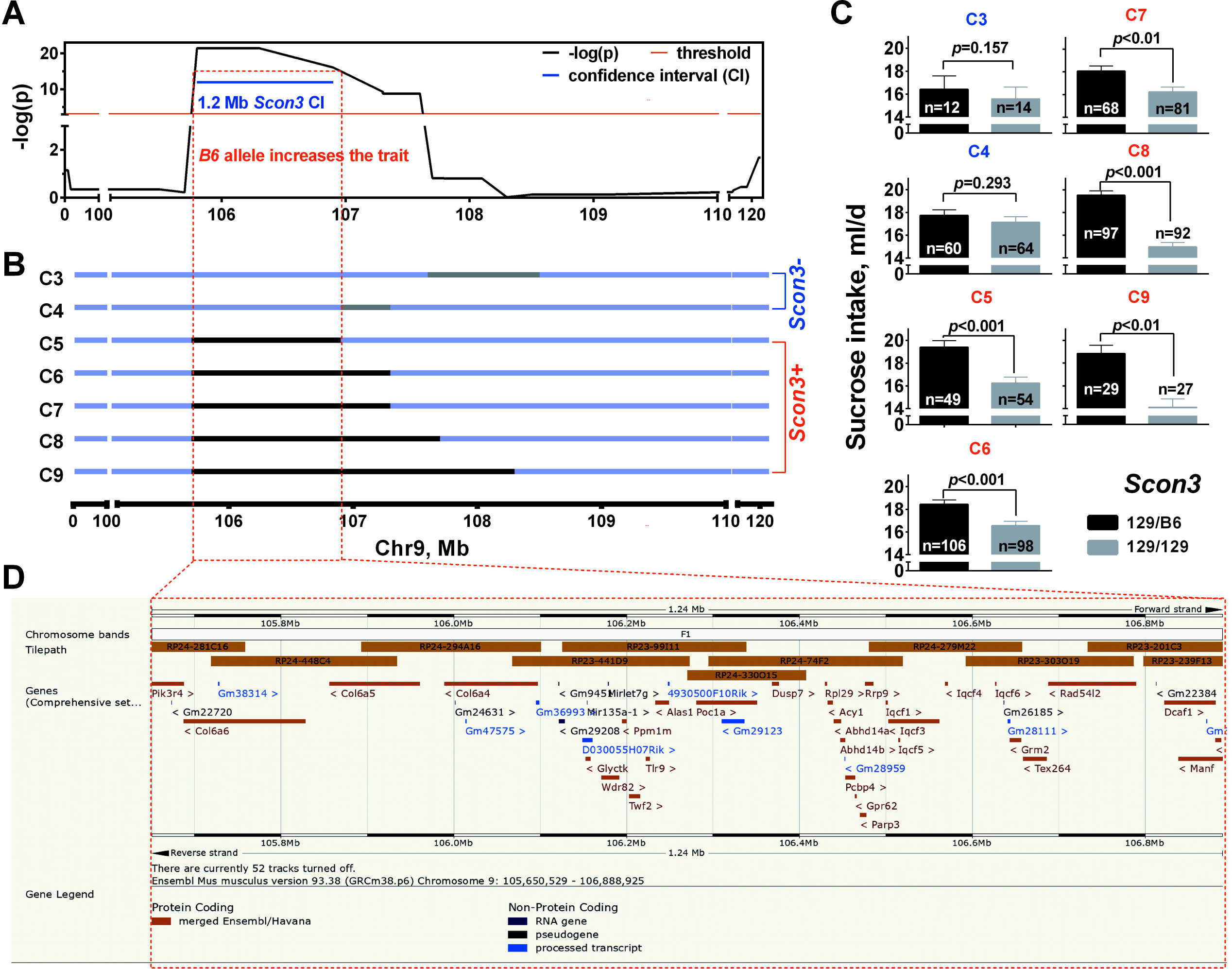
High-resolution mapping of the *Scon3* locus in congenic substrains. (**A**) For average daily sucrose intake, we computed trait-marker associations using a general linear model among all 129.B6-*Scon3* congenic mice grouped by genotype at each marker. The x-axis gives marker positions in Mb on chromosome 9 (Chr9); for other details see **Figure 3**. (**B**) Donor region of each congenic strain: black bars, strains that retained the *Scon3* locus; gray bars, strains that lost the locus; blue bars, region contributed by the host strain. (**C**) Strain comparisons with sample size (*n*) of each genotype within each strain. C3 and C4, *Scon3*-negative strains; C5–C9, *Scon3*-positive strains that share a common region (red dashed rectangle in **A and B**: 1.2-Mb donor region between 105.7 and 106.9 Mb, from *rs33653996* to *rs3023231*) that the *Scon3*-negative strains (C3 and C4) do not. (**D**) The 1.2 Mb *Scon3* critical region contains X noncoding RNA genes, Y protein-coding genes, Z pseudogenes, and N processed transcripts (Ensemble Mouse Genome Brower @GRCm38.p5 (Hubbard, Aken et al. 2007)).

#### Scon3 and Scon4 interaction in double congenics

To confirm epistatic interaction between *Scon3* and *Scon4* detected in F_2_ (**Figures 1B, C)**, we intercrossed *Scon3* and *Scon4* single congenics to produce mice with B6 donor alleles of both QTLs segregated (double congenic, **Figure 2B**). To achieve this, we used a homozygous *Scon4* congenic male (B6/B6; N_5_F_2_) mated with heterozygous *Scon3* congenic females (B6/129; N^7^). Consequently, all offspring were heterozygous (129/B6) for *Scon4* but had segregating (129/B6 heterozygotes and 129/129 homozygotes) *Scon3* genotypes (**Figures 2G, H**). The double congenic mice from this substrain that had B6 alleles at both loci (129/B6 genotype for *Scon3* and *Scon4*) drank three times more sucrose than their littermates with genotypes 129/B6 of *Scon4* and 129/129 of *Scon3* (Cohen’s *D*=5.0); mice with only one B6 allele drank much less (Cohen’s *D*: *Scon4*, 0.9; *Scon3*, 0.8; **Figures 2H, I**). For the genotypes for *Scon4* and *Scon3*, refer to **Figure 2G**, except for congenic strains C2, C3 and C4, which we recoded as 129/129 because these strains did not retain the QTLs (**Figures 3B, D**, and **Figures 5B, C**). Thus, mice with three genotypes were included in the pooled congenic analyses: 129.B6-*Scon4* (129/B6 of *Scon4*, 129/129 of *Scon3*; mice from substrain C1), 129.B6-*Scon3* (129/129 of *Scon4*, 129/B6 of *Scon3*; mice from substrains C5-C9), and mice without B6 allele (129/129 of *Scon4*, 129/129 of *Scon3;* mice from half of substrains C1, C5-C9 and all of substrains C2-C4). A single model encompassing all the congenic data produced results very similar to those of the F_2_ and consomics: mice differed in voluntary sucrose intake depending on the combination of *Scon3* and *Scon4* genotypes [*Scon3*: *F*(1,989)=109.7, *p*<0.00001; *Scon4*: *F*(1, 989)=8.65, *p*<0.005; *Scon3* × *Scon4*: *F*(1, 989)=40.2, *p*<0.00001; **Figure 2J**].

### Characterization of the *Scon3 and Scon4* QTL phenotypes

#### Electrophysiology

No statistically significant differences from 129 mice were found for 129.B6-*Scon3* heterozygotes or 129.B6-*Scon4* heterozygotes for any of the taste stimuli tested (100, 300 and 1000 mM sucrose; 20 mM quinine; 100 mM NaCl; 10 mM HCl; 1000 mM glycine; 1000 maltose; 1000 mM fructose; 2, 6, and 20 mM saccharin; 100, 300, and 1000 mM MSG; and 3% nd 10% ethanol; *p*>0.05, Bonferroni correction for multiple tests; **S6 Figure**, S8 Table)

In the following characterization the *Scon4* QTL effect, we compared homozygous *Scon4* congenics (B6/B6) with 129 host inbred mice (129/129).

#### Two-bottle preference tests

The *Scon4* genotype affected the consumption of 300 and 600 mM sucrose, 600 mM glucose, and 600 mM fructose, but it had no effect at other, mostly lower concentrations of these sugars (**Figure 6**). Likewise, *Scon4* congenic and 129 inbred mice did not differ in their consumption of 20 mM saccharin, 30 mM glycine, or 10% ethanol (**S7A, B, F Figures**) or of 4.6% soybean oil, 6% cornstarch, or 10.5% maltodextrin (**S8A-C Figures**).However, *Scon4* congenic mice had lower intakes of 300 mM NaCl, 300 mM MSG, and 3% ethanol and lower 3% ethanol preference scores compared with 129 inbred mice (**S7C-E Figures**).

**Figure 6.**
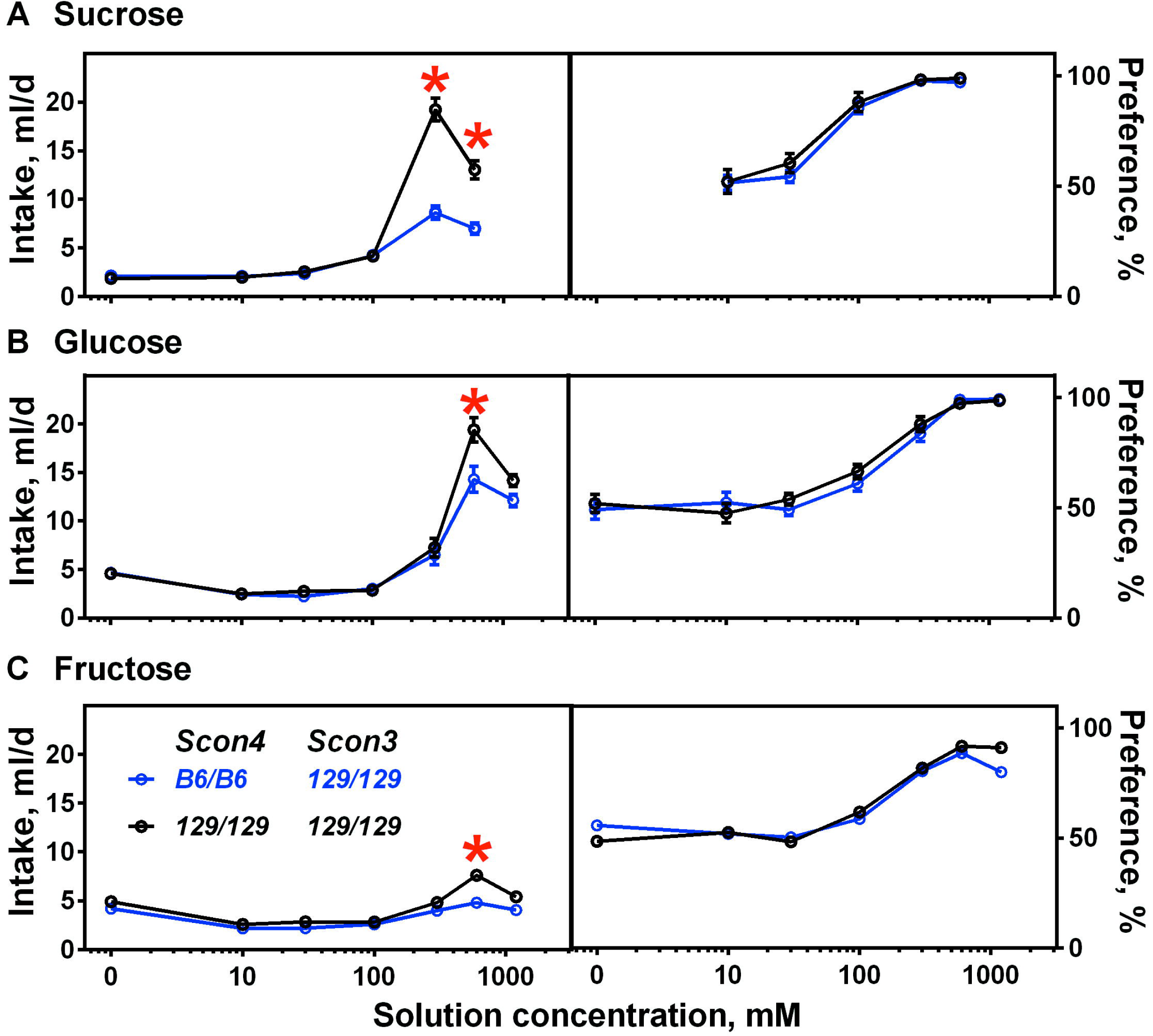
The effects of *Scon4* are largest for the more concentrated sugar solutions Average daily intake (left panels) and preference (right panels) for sucrose. (**A**), glucose (**B**), and fructose (**C**) by homozygous 129.B6-*Scon4* congenic mice (C1; for sucrose, *N*=19; for glucose, *N*=10; for fructose, *N*=10) and host inbred 129 mice (all *N*=10) in two-bottle 48-hr preference tests. Error bars show SEM. \**p*<0.05, significant difference between the congenic and host inbred mice (post hoc tests). Blue, genotypes for *Scon4* and *Scon3*; black, control 129 inbred mice.

#### Body composition analysis

We used three methods to study body composition, MR, DEXA, and necropsy, all of which indicated that the *Scon4* genotype had no significant effect on body composition (MR, **S9A Figure**; DEXA, **S9B Figure**; necropsy, **S10 Figure**). From the necropsy data, we learned that mice of different genotypes had similar organ weights, except for the subscapular adipose depot (von Mering, Jensen et al. 2005) and heart (**S10 Figure**). Drinking sucrose voluntarily for 4-12 days did not result in significant differences in body weight gain among mice of different *Scon3* and *Scon4* genotypes (**S11 Figure**).

#### Metabolism

Homozygous *Scon4* mice differed in their metabolic gas exchange compared with inbred host controls, especially in early adulthood. Specifically, mice from the C1 strain (B6/B6) consumed less oxygen, generated less heat (kcal/kg lean), and burned proportionately more calories as fat than carbohydrate compared with inbred 129 mice (129/129; **Figures 7C-F**). However, at both ages tested, these homozygous *Scon4* mice ate the same amount of food (g/24 hr), drank the same amount of water (mL/24 hr), and were as active (beam breaks/hr) as inbred 129 mice (**Figures 7A, B, G)**.

**Figure 7.**
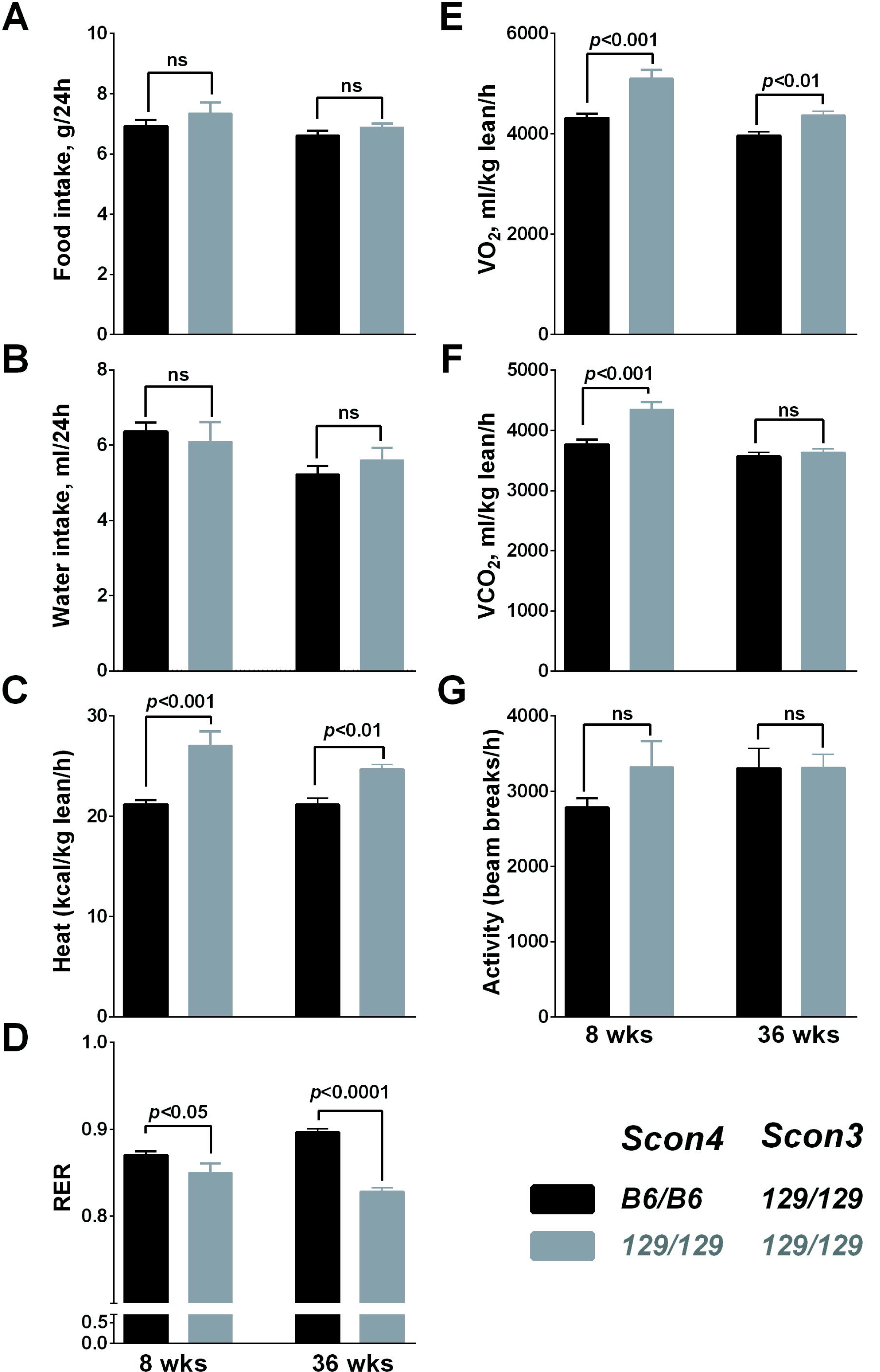
The *Scon4* genotype affects fuel oxidation but not food intake. Metabolic assessments in *Scon4* congenic mice (black bars; *N*=17; C1) and control 129 inbred mice (gray bars; *N*=8) at 8 and 36 weeks of age. (**A**) Food intake: 8 weeks, *t*_(1,23)_=1.11, *p*=0.282; 36 weeks, *t*_(1,23)_=1.10, *p*=0.284. (**B**) Water intake: 8 weeks, *t*(_1,23)_=0.54, *p*=0.597; 36 weeks, *t*_(1,23)_=0.96, *p*=0.348. (**C**) Heat production: 8 weeks, *t*_(1,23)_=5.22, *p*<0.001; 36 weeks, *t*_(1,23)_=3.49, *p*<0.01. (**D**) Respiratory exchange ratio (RER): 8 weeks, *t*_(1,23)_=5.22, *p*<0.001; 36 weeks, *t*_(1,23)_=3.49, *p*<0.01. (**E**) Oxygen consumption (VO_2_): 8 weeks, *t*_(1,23)_=4.71, *p*<0.001; 36 weeks, *t*_(1,23)_=3.16, *p*<0.01. (**F**) Carbon dioxide (VCO_2)_: 8 weeks, *t*_(1,23)_=4.03, *p*<0.001; 36 weeks, *t*_(1,23)_=0.53, *p*=0.598. (**G**) Activity over the entire 24-hr light/dark cycle: 8 weeks, *t*_(1,23)_=1.85, *p*=0.077; 36 weeks, *t*_(1,23)_=0.00, *p*=0.999. Data are mean ± SE.

#### Glucose tolerance test

Homozygous *Scon4* mice did not differ from inbred 129 mice in blood glucose or insulin in response to exogenous glucose, by either IP or IG administration, whether tested at 8 weeks or at 36 weeks (**S12A, B Figures**).

#### Insulin tolerance test

After injection with insulin in 8-week-old mice, a significant genotype × time interaction was detected, with *Scon4* congenic mice tending to have higher insulin sensitivity (lower blood glucose) than 129 inbred mice, with post hoc tests detecting significant difference at 90 min (**S12C Figure**, left panel). The same mice at 36 weeks showed no effects of genotype at all and no interaction of genotype with time (**S12C Figure**, right panel).

### Candidate genes

We identified and investigated genes in the *Scon3* and *Scon4* regions defined by the congenic mapping results (*Scon3*, 9:105650529-106888925; *Scon4*, 1:1-25377067). *Scon3* harbors 55 genes and *Scon4* harbors 193 genes (S9 Table), of which 49 are protein-coding genes [21] and 199 are other gene types, such as long noncoding RNA (lncRNA) genes. Among 41 of the protein-coding genes, there were 168 relevant missense and stop codon gain/loss variants (S9 Table). We grouped the biological processes for these genes into categories (**S13A Figure**) and found that 31 of these known genes are involved in carbohydrate metabolism (S9 Table). We identified pairs with the highest association confidence scores and containing one gene from the *Scon4* region and another from the *Scon3* region. For example, *Rb1cc1* (RB1-inducible coiled-coil 1) from *Scon4* and *Pik3r4* (phosphatidylinositol 3 kinase, regulatory subunit, polypeptide 4, p150) from *Scon3* have an interaction confidence score > 0.9 (**S13B Figure**). The prioritization of these two genes ranks first and eleventh based on the similarity to other genes involved in carbohydrate metabolism, and one member of the pair (*Pik3r4*) has a missense variation (S9 Table). Additional gene pairs also had protein-protein association scores > 0.9: *Oprk1*-*Grm2* (opiate and glutamine receptors), *Grm2*-*Npbwr1* (glutamine and neuropeptide receptors), and *Rpl29*-*Rpl*7 (ribosomal proteins) (S10 Table).

Gene expression profiling at three mouse tissue sites (striatum, brown and white fat) revealed a significantly differential expression of 20 candidate genes (*2610203C22Rik, Abhd14a, Adhfe1, Col6a4, Col6a5, Dusp7, Eya1, Glyctk, Iqcf1, Kcnq5, Mcm3, Paqr8, Parp3, Poc1a, Prex2, Sgk3, Snhg6, Sox17, St18, Ube2w*) within *Scon3* and *Scon4* (**Figure 8;** S11 Table) between 129 and B6 inbred mouse strains (twofold change with FDR<0.05).

**Figure 8.**
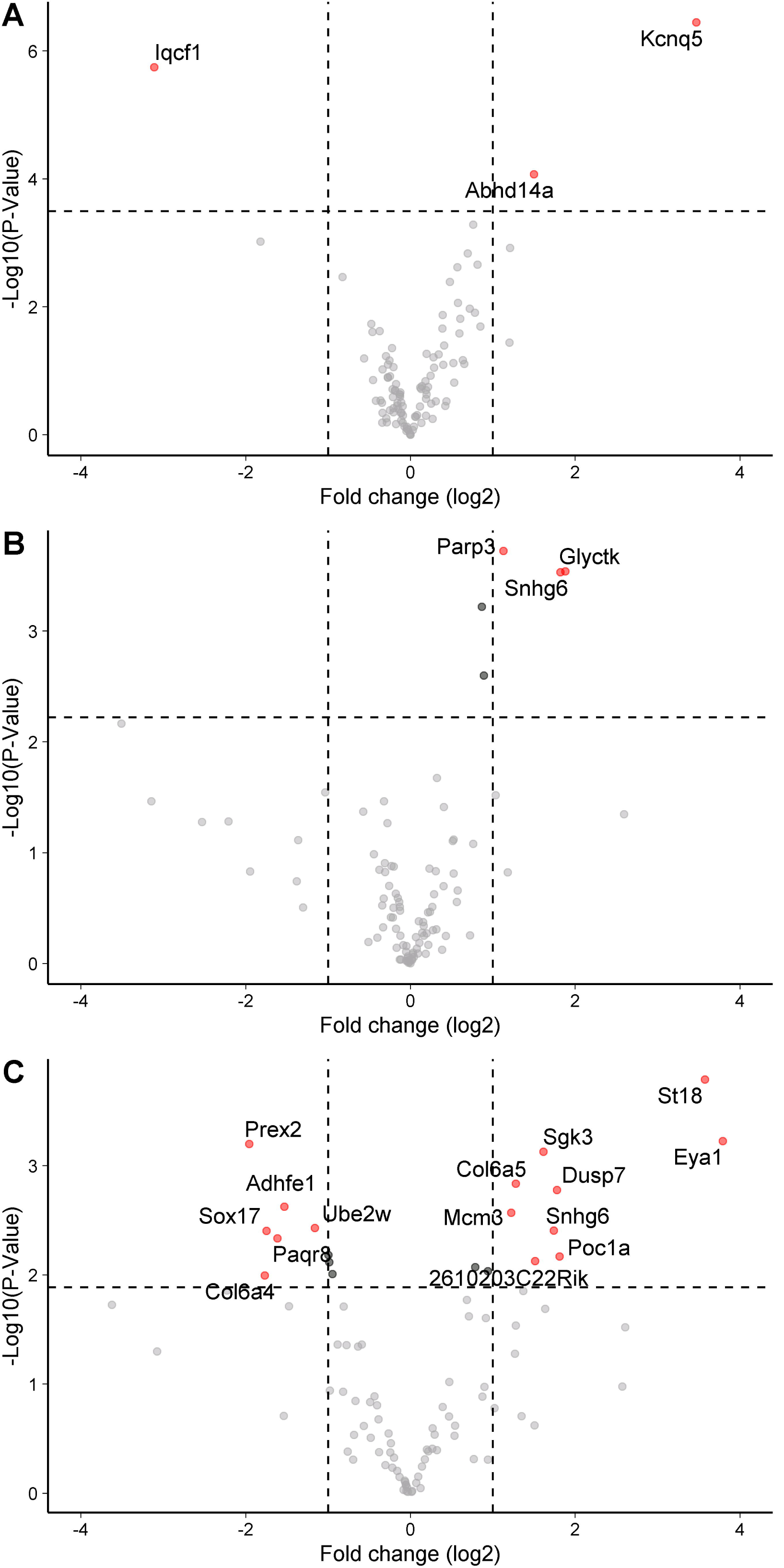
Gene expression profiling shows differential expression of candidate genes within *Scon3* and *Scon4* regions between 129 and B6 inbred mouse strains. Data are from microarray-based gene expression analysis for mouse stratum (**A**) and bulk RNASeq analyses for brown (**B**) and white (**C**) fat. Red dots show differentially expressed genes with twofold changes between two inbred strains. The horizontal dash lines show the corrected significance threshold (FDR=0.05).

## Discussion

We identified *Scon3* and *Scon4*, two small regions of the mouse genome containing variations that affect the voluntary intake of concentrated sucrose solution, which emerged when influential variation in the *Tas1r3* gene was removed. We used B6 and 129 mice for these experiments because of the marked differences in sucrose consumption between the two strains (Bachmanov, Tordoff et al. 2001) and because we were interested in learning more about the region on chromosome 9 identified in an earlier genome scan (see the companion paper (Lin 2020)). The utility of these mice was confirmed here by the initial *Tas1r3* -independent intercross results, which confirmed the striking influence of a locus on Chr9 that interacts with a separate region on Chr1.

We are unable to draw firm conclusions about the *Scon4* locus because we did not test all taste solutions in both naive and experienced mice (S12 Table). The parental inbred strains are susceptible to develop learned flavor preferences and thus prior experience with nutrients changes their response to taste solutions (Sclafani 2006), so we cannot directly compare the results of tests conducted with naive mice with those conducted on mice with prior experience. Thus, while we can be sure that naive mice differ in sucrose intake by *Scon4* genotype, but we cannot be sure that there is no effect of genotype on starch intake in naive mice because mice drank another nonsweet but preferred solution beforehand (soybean oil).

We found that the *Scon4* genotype was related to whole-body fuel use. Under basal conditions, mice with genotypes of *Scon4* that decreased voluntary sucrose consumption also oxidized more carbohydrate and less fat than did mice with opposing genotypes. These genetic effects on voluntary sucrose intake and carbohydrate oxidation do not seem to arise from the earliest steps of intestinal absorption because there were few or no differences by genotype in the outcomes of glucose and insulin tolerance tests. However, we cannot rule out differential stimulation of postoral nutrient detectors that may partition nutrient oxidation and storage.

Peripheral taste sensitivity might be a contributor to the differences in voluntary sucrose consumption, but four points make this explanation unlikely: (a) genotype affects voluntary intake at high but not low sucrose concentrations (the ability to detect sweet taste is more important at lower concentrations), (b) genotype effects studied here were absent for peripheral nerve responses, (c) these small effects were in the opposite direction from those of another genetic model of voluntary sucrose intake (Inoue, Reed et al. 2004), and (d) we found no relationship between *Scon4* or *Scon3* in a genome scan using taste nerve responses as a trait (see the companion paper (Lin 2020)). However, we cannot rule out a peripheral effect entirely, given the small sample size of mice tested for peripheral neural responses to sucrose.

There are at least two ways to approach the evaluation of genes within the critical intervals for their effect on voluntary sucrose consumption. The first method is to examine all genes within the regions to determine if there are known connections between those genes and voluntary sucrose consumption (narrowly) and metabolism (more broadly). No genes were linked to voluntary sucrose consumption in the databases surveyed, but many were part of carbohydrate and fat metabolism pathways, including *Pik3r4* (Nemazanyy, Blaauw et al. 2013), *Alas1* (Simcox, Mitchell et al. 2015), *Acy1* (Alexandre-Gouabau, Bailly et al. 2012), *Cops5* (Tian, Peng et al. 2010), *Atp6v1h* (Olsson, Yang et al. 2011, Baumeier, Kaiser et al. 2015), *Tlr9* (Lewis and Cobb 2010), *Lypla1* (Hacia, Fan et al. 1999, Mootha, Bunkenborg et al. 2003), *Pkhd1* (Kaimori, Nagasawa et al. 2007), and *Gdap1* (Lopez Del Amo, Palomino-Schatzlein et al. 2017). We further reasoned that the interaction of the *Scon3* and *Scon4* regions indicated that causal genes might be part of the same pathway. That approach led us to identify particular gene pairs that jointly contribute to a shared function (e.g., *Pik3r4* and *Rb1cc1*); however, their involvement in voluntary sucrose intake awaits direct tests.

The second approach is to find genes or gene pathways that have a similar constellation of traits, with the idea that the genes in the region could be unknown members of the known pathway. Using this second method, we learned that many of the traits of the *Scon4* and *Scon3* mice studied here are similar to mice with knockout or overexpression of the *FGF21* gene. These traits include increased intake of sugar and the lack of large effects on peripheral taste nerves and body weight or fatness (von Holstein-Rathlou, BonDurant et al. 2016). A next step would be to evaluate the genes in the region experimentally for their roles in the *FGF21* signaling pathway. We note that two proteins that regulate this signaling pathway, *Prdm14* (Grabole, Tischler et al. 2013) and *Sulf1* (Buresh, Kuslak et al. 2010), are located in the *Scon4* region. *SGLT1* and *KATP*, as an alternate potential sucrose transduction mechanism (Yee, Sukumaran et al. 2011), are located on different chromosomes from *Scon3* and *Scon4*, and it is unlikely there is a peripheral taste effect of *Scon3* and *Scon4*, so they can be ruled out as candidates.

Through our use of consomic mouse strains, we indirectly mapped other genes for taste-related traits. Consomic strains contain an entire introgressed chromosome; thus, there is a greater potential to capture more causal genes and their variants compared with the congenic strains, which have smaller introgressed regions. As expected, our consomic mapping results discovered new regions of interest distinct from the *Scon* loci and also confirmed many previously reported regions that contain variation that affects intake of several other taste compounds by mice (Phillips, Crabbe et al. 1994, Tarantino, McClearn et al. 1998, Vadasz, Saito et al. 2000, Tordoff, Reed et al. 2008).

This study was conducted in mice, but what we learned has connections to human health. Although the contribution of sucrose, especially in beverages, to the ills of the modern diet remains controversial, the benefits of understanding the biological causes of excessive sugar consumption are likely to be helpful. An earlier example of this type of benefit comes from the discovery of a subunit of the mouse sweet receptor, T1R3 (Bachmanov, Li et al. 2001, Kitagawa, Kusakabe et al. 2001, Max, Shanker et al. 2001, Montmayeur, Liberles et al. 2001, Nelson, Hoon et al. 2001, Sainz, Korley et al. 2001). This discovery was useful in understanding taste biology, but it is now apparent that this sugar-sensing receptor has many nongustatory functions, including insulin regulation (Geraedts, Takahashi et al. 2012) and adipogenesis (Simon, Parlee et al. 2013). Likewise, we hope this study provides insights toward understanding the appetition for sugar and that this knowledge can be harnessed to help prevent the negative effects of excessive sugar consumption by understanding and allaying the human hunger for sweetness.

## Acknowledgments

We gratefully acknowledge Maria L. Theodorides, Zakiyyah Smith, Mauricio Avigdor, and Amy Colihan for assistance with animal breeding. We also acknowledge Richard Copeland and the consistent high-quality assistance of the animal care staff at the Monell Chemical Senses Center and thank them for their service. Matt Kirkey assisted with genotyping the congenic mice. Yutaka Ishiwatari assisted with genotyping markers. Longhui Chen assisted with metabolic experiments. Anthony Sclafani made constructive comments on an earlier draft of the manuscript.

## Ethics approval

All animal study procedures were approved by the Monell Chemical Senses Center Institutional Care and Use Committee.

## Consent to participate

Not applicable.

## Consent for publication

Not applicable

**Availability of data and material:**

## Code availability

**Not applicable**.

**Accession IDs:**

## Supplemental Figures

**S1 Figure. Correlations between sucrose intake (ml/d) and phenotype within each genetic mapping population** Data are for body weight (BW; g) by sex (M, F), average daily habitual water intake (Wat; ml/d, M+F combined), and age (weeks, M+F combined). For sample sizes, see S1 Table.

**S2 Figure. F**_**2**_ **mice (*N*=279) showed genotype effects of the *Scon3 and Scon4* loci on intake or preference for taste solutions** Group means that do not differ either have no letter subscript or share a letter subscript in common. A=B6/B6, H=129/B6, B=129/129.

**S3 Figure. No *Scon3* or *Scon4* genotype effects on average daily water intake in F**_**2**_ **(A, B), congenic (C, D), or backcross (E) mice** For sample sizes, see S1 Table. A=B6/B6; H=129/B6; B=129/129. Average daily water intake was analyzed using general linear model with genotype and sex as fixed factors and body weight as covariate. Within each panel, groups that do not share a common subscript letter differ significantly (post hoc tests).

**S4 Figure. Heat map of Pearson correlations for taste solution intakes and preference scores of the F**_**2**_ **mice (*N*=279)**

**S5 Figure. Chromosomes 1 and 9 harbor QTLs for the consumption of other taste solutions**. A significant difference in intake and/or preference of a solution between a consomic strain and host inbred 129 or B6 strain indicates presence of a QTL on this chromosome for the taste solution (*p*<0.05). For intakes, we used a general linear model, with strain as fixed factor and habitual water intake as a covariate, followed by post hoc test. For preference scores, we used *t*-tests.

**S6 Figure. Both *Scon3* and *Scon4* showed no genotype effect on the responses of the chorda tympani gustatory nerve to lingual application of taste stimuli** We compared the heterozygous 129.B6-*Scon4* congenic mice (C1; B6/129; *N*=5) or homozygous 129.B6-*Scon3* congenic mice (C7: B6/B6; *N*=5) and host inbred 129 mice (129/129; *N*=5) using Mann-Whitney *U*-tests for individual solutions. We observed no significant genotype differences (*p*>0.05) in responses of the chorda tympani gustatory nerve for the taste stimuli (see S8 Table). Values are medians ± median absolute deviations.

**S7 Figure. Effects of the *Scon4* genotype on intakes and preference for other taste compounds** Data are for 20 mM saccharin (**A**), 30 mM glycine (**B**), 300 mM NaCl (**C**), 300 mM MSG (**D**), 3% ethanol (**E**), and 10% ethanol (**F**) by homozygous 129.B6-*Scon4* congenics (C1: B6/B6) and host inbred 129 mice (129/129) in two-bottle tests.

**S8 Figure. No effect of the *Scon4* genotype on intakes and preferences for other caloric compounds** Data are for 4.6% soybean oil (**A**), 6% cornstarch (**B**), or 10.5% maltodextrin (**C**) by homozygous 129.B6-*Scon4* congenics (C1: B6/B6; *N*=19) and host inbred 129 mice (129/129; *N*=10) in two-bottle tests.

**S9 Figure. No effect of the *Scon4* genotype on mouse body composition at 8 or 36 weeks of age** Mouse body composition as measured by magnetic resonance (**A**; 8 weeks: *N*=32, males=15, females=17; 36 weeks: *N*=22, males=9, females=13) or by DEXA (*N*=26, males=12, females=14): homozygous *Scon4* congenics (C1; B6/B6) vs host inbred 129 controls (129/129). BMC=bone mineral content (g); BMD=bone mineral density. Groups that do not share a common subscript letter differ by post hoc test.

**S10 Figure. Effect of *Scon4* genotype on organ and fat weight of mice dissected at 180 days of age** No effect was observed, except for subscapular adipose depot (*F*_(1,23)_=18.5, *p*<0.001) and heart (*F*_(1,23)_=22.5, *p*<0.0001). Homozygous *Scon4* congenics (C1; B6/B6; males=7, females=12) vs host inbred 129 controls (129/129; males=5, females=2). Groups that do not share a common subscript letter differ by post hoc tests.

**S11 Figure. No effect of *Scon4* genotype on body weight changes when mice were fed chow and drank sweetened water** Water was sweetened with sucrose (**A-F**), glucose (**G**), or fructose (**H**) solution. Mice drank voluntarily for 4-12 days. Data are for F_2_ mice (**A, B**; *N*=279), consomic mice (**C-E**; 129.B6-Chr1 and 129 controls: *N*=105; 129.B6-Chr9 and 129 controls: *N*=87; B6.129-Chr9 and B6 controls: *N*=86), or congenic mice (**F-H**; sucrose: *N*=105; glucose: *N*=20; fructose: *N*=20). Groups that do not share a common subscript letter differ by post hoc test. A=B6/B6; H=129/B6; B=129/129.

**S12 Figure. No effect of *Scon4* genotype on tolerance to glucose or insulin** Glucose was administered intraperitoneally (IP GTT; **A**) or intragastrically (IG GTT; **B**) or insulin was administered (**C**) in homozygous *Scon4* congenic mice (black squares; *N*=21) and the host inbred control mice (gray circles; *N*=12) at 8 and 36 weeks of age. Data are mean ± SEM of genotype; significance of the genotype effect was evaluated by post hoc tests. \**p*<0.05. The statistics are as follows: **IP**, 8 weeks, genotype *F*_(1,30)_=1.52, *p*=0.227; time *F*_(7,210)=_73.5, *p*<0.0001, genotype × time interaction *F*_(7,210)_=1.70, *p*=0.111; 36 weeks, genotype *F*_(1,24)_=0.19, *p*=0.665; time *F*_(7,168)_=107.9, *p*<0.0001; genotype × time interaction *F*_(7,168)_=1.70, *p*=0.112. **IG**, 8 weeks, genotype *F*_(1,31)_=1.95, *p*=0.173; time *F*_(7, 217)_=69.4, *p*<0.0001; genotype × time interaction (*F*_(7,217)_=0.18, *p*=0.989); 36 weeks, genotype *F*_(1,24)_=0.065, *p*=0.800; time *F*_(7,168)_=62.2, *p*<0.0001; genotype × time interaction *F*_(7, 168)_=0.53, *p*=0.813. **ITT**, 8 weeks, genotype *F*_(1,28)_=8.68, *p*=0.006; time *F*_(5, 140)_=101.2, *p*<0.0001; genotype × time interaction *F*_(5,140)_=3.70, *p*=0.003; 36 weeks, genotype *F*_(1,23)_=1.012, *p*=0.325; time *F*_(5,115)_=80.1, *p*<0.0001; genotype × time interaction *F*_(5,115)_=1.34, *p*=0.253.

**S13 Figure. Gene ontology analysis and protein-protein associations for genes within the *Scon3* and *Scon4* regions** (**A**) Gene ontology analysis of the biological process for genes within the *Scon4* and *Scon3* regions. The Ensembl mouse gene identifier IDs (*N*=195; no Ensembl gene identifier IDs were available for the 45 unknown genes) were batch-uploaded into the Panther database for *Mus musculus*, and in total, 92 genes and 72 hits (gene IDs matched in the public database) were processed. (**B**) Protein-protein associations for genes within *Scon4* and *Scon3* (S10 Table) were identified by searching protein names using information from high-throughput experimental data, mining of literature databases, and predictions based on genomic context analysis.

