## Supplemental Table 2 for "Genetic controls of mouse *Tas1r3*-independent sucrose intake"

**S2 Table.** The consomic strains used in this study

| **Official strain symbol^a^** | **Abbreviation** | **MMRRC ID^b^** | **JAX ID^b^** |
| --- | --- | --- | --- |
| 129P3/J-Chr 1^C57BL6ByJ^/MonMmjax | 129-Chr1^129^ | 36684 | 018675 |
| 129P3/J-Chr 9^C57B6/ByJ^/ MonMmjax | 129-Chr9^129^ | 36687 | 018678 |
| C57BL/6ByJ-Chr 9^129P3/J^/ MonMmjax | B6-Chr9^129^ | 36690 | 018681 |

^a^ “Mon” within mouse strain name is a laboratory code for the Monell Chemical Senses Center issued by the Institute for Laboratory Animal Research (ILAR; http://dels.nas.edu/ilar_n/ilarhome/labcode.shtml).

^b^Identification numbers (ID) are for strains available from the Mutant Mouse Regional Resource Center (MMRRC; https://www.mmrrc.org) and The Jackson Laboratory (JAX; http://jaxmice.jax.org).
