## Supplemental Table 12 for "Genetic controls of mouse *Tas1r3*-independent sucrose intake"

**S12 Table.** Summary of sucrose intake and genotype of *Scon4* and *Scon3* of the experimental mouse crosses

| Population | *N* | Type | *Scon4*  genotype | *Scon3* genotype | Sucrose intake  (mean ± SE, ml/day) |
| --- | --- | --- | --- | --- | --- |
| F_2_^a^ | 20 | B6 x 129.B6^Sac^ | B6/B6 | B6/B6 | 27.4 ± 0.9 |
| F_2_^a^ | 28 | B6 x 129.B6^Sac^ | 129/B6 | B6/B6 | 26.9 ± 0.8 |
| F_2_^a^ | 18 | B6 x 129.B6^Sac^ | 129/129 | B6/B6 | 27.5 ± 1.0 |
| F_2_^a^ | 34 | B6 x 129.B6^Sac^ | B6/B6 | 129/B6 | 27.1 ± 0.7 |
| F_2_^a^ | 69 | B6 x 129.B6^Sac^ | 129/B6 | 129/B6 | 26.1 ± 0.5 |
| F_2_^a^ | 28 | B6 x 129.B6^Sac^ | 129/129 | 129/B6 | 26.4 ± 0.7 |
| F_2_^a^ | 13 | B6 x 129.B6^Sac^ | B6/B6 | 129/129 | 14.3 ± 1.2 |
| F_2_^a^ | 28 | B6 x 129.B6^Sac^ | 129/B6 | 129/129 | 17.7 ± 0.8 |
| F_2_^a^ | 23 | B6 x 129.B6^Sac^ | 129/129 | 129/129 | 23.1 ± 0.9 |
| Inbred | 43 | B6 | B6/B6 | B6/B6 | 23.1 ± 0.5 |
| Inbred | 64 | 129 | 129/129 | 129/129 | 18.1 ± 1.5 |
| Backcross | 157 | F1 x 129, N3-N6 | Not tested | B6/129 | 20.7 ± 0.3 |
| Backcross | 280 | F1 x 129, N3-N6 | Not tested | 129/129 | 16.9 ± 0.3 |
| Consomics | 39 | 129.B6-Chr1 | B6/B6 | 129/129 | 11.8 ± 0.7 |
| Consomics | 28 | 129.B6-Chr9 | 129/129 | B6/B6 | 23.4 ± 0.9 |
| Consomics | 43 | B6.129-Chr9 | B6/B6 | 129/129 | 14.6 ± 0.5 |
| Congenics | 20 | 129.B6-Scon4, hetero | 129/B6 | 129/129 | 14.8 ± 0.9 |
| Consomics | 15 | Littermates^b^ | 129/129 | 129/129 | 18.6 ± 1.1 |
| Consomics | 40 | 129.B6-Scon4, homo | B6/B6 | 129/129 | 8.2 ± 0.6 |
| Consomics | 348 | 129.B6-Scon3, hetero | 129/129 | B6/129 | 18.8 ± 0.2 |
| Consomics | 352 | Littermates^c^ | 129/129 | 129/129 | 15.8 ± 0.2 |
| Consomics | 5 | 129.B6-Scon3, homo | 129/129 | B6/B6 | Not tested |
| Consomics | 14 | 129.B6-double | 129/B6 | 129/B6 | 23.1 ± 0.7 |
| Consomics | 12 | Littermates^d^ | 129/B6 | 129/129 | 8.3 ± 0.9 |

^a^F_2_ mice drank more sucrose regardless of genotype likely because all F_2_ mice have the B6 allele at *Tas1r3* that increases sucrose intake and because we tested with 50 mM sucrose for 4 consecutive days before testing with 300 mM sucrose.

Littermates from the 129.B6-*Scon4* strain but *Scon3* retains the 129 allele.

^c^Littermates from the 129.B6-*Scon3* strain with the 129 allele at the *Scon3* locus.

^d^Littermates from the 129.B6-double strain; only *Scon4* retains the B6 allele.
